# Anatomical Plasticity of the Distal Forelimb Projection of the Ventral Premotor Cortex Four weeks After Primary Motor Cortex Injury

**DOI:** 10.1101/2020.06.12.148494

**Authors:** David W. McNeal, Scott Barbay, Shawn B. Frost, Michael Taylor, David J. Guggenmos, Randolph J. Nudo

## Abstract

Brain injury affecting the isocortical frontal cortex is a common pathological occurrence. Many patients report severe deficits to functions of daily living. However, there is a variable degree of motor recovery that occurs with some individuals recovering astounding degrees of motor recovery while others have not. This variability has led researchers into investigating the possible mechanisms for this variability. Recently, several non-human primate studies have shed light on the possibility of spared, ipsilesional motor area taken over the lost function to the damaged cortex. Unfortunately, these studies have focused on long-term adaption ranging from 5months to one year post injury. In this present study, we are the first use rigorous stereological quantification to show that significant neuroplastic changes in the form of changes to neuroanatomical connections between distant cortical area occurs at a very early time point of 4 weeks post injury. Much like the Dancause study in 2005, we found that ishemic damage to the distal forelimb area (DFL) of the primary motor cortex (M1) induced plastic changes between the DFL of the ventral premotor cortex (PMv) and area 1/2 of the somatosensory cortex. Indeed, we found a nearly 2 fold increase in the number of boutons between PMV and area 1/2. Additionally, labeled fibers from PMv change direction from their normal termination within M1 and traveled in a ventral posterior direction toward the somatosensory cortex. Also of interest, several labeled fibers actually traveled through the glial scar of M1 toward the somatosensory cortex. These data demonstrate that a massive neuroplastic response has occurred following an ischemic insult to the DFL of M1. These data may suggest that the brain may be undergoing an attempt to re-establish a degree of motor and or sensory control to compensate for the lost function due to the injury.

## Introduction

Stroke is defined as a major episode of interruption of blood flow to the cerebral cortex, which leads to ischemic damage, and death of glial cells and cortical neurons. When the ischemic damage affects glial cells and neurons comprising the arm representation of the primary motor cortex (M1), there is a presentation of a distinct contralateral paresis of proximal and distal movements of the upper extremity with distal movements exhibiting a more prominent and severe paresis (Chollet et al. 1991). As one would expect, this paresis of the distal extremities becomes a major hindrance to accomplishing even the most basic and self-reliant functions of daily living for stroke survivors. Indeed, nearly 30% of all stroke survivors report such severe deficits in the distal musculature of the hand they are classified with a major disability thereby making stroke a major long-term health issue in the United States (American Heart Association, 2013). However, some stroke survivors demonstrate a degree of spontaneous motor recovery of the upper extremity allowing for some restoration of dexterity and elevated levels of daily functioning although this recovery is highly variable among patients (Duncan et al. 2000; Stinear 2010). What drives this recovery in some individuals and not in others has become a major focus in clinical and neuroscience research.

Recognition of the limited knowledge into the underlying mechanisms of recovery of lost motor function after stroke has spurred increased interest in human and animal post-injury cortical mechanisms over the past 15 years (Frackowiak et al. 1991; Weiller et al. 1992; Morecraft and Van Hoesen 1993; Nudo and Milliken 1996; Nudo 1997; Feydy et al. 2002; Nudo et al. 2003; Ward et al. 2003; Dancause, Barbay, Frost, Plautz, Popescu, et al. 2006; Pizzimenti et al. 2007; Eisner-Janowicz et al. 2008; McNeal et al. 2010; Darling, Pizzimenti, and Morecraft 2011; Nudo and McNeal 2013). Importantly, based upon human functional imaging studies showing metabolic increases in “spared” motor areas of the cerebral cortex residing on the same side of the cortex as the lesion, researchers have begun to suspect that these areas may play a significant role in motor recovery (Frackowiak et al. 1991; Weiller et al. 1993; Picard and Strick 2001; Ward et al. 2003). In conjunction with these human imaging studies, animal studies have investigated neurophysiological investigations into the cortical mechanisms of recovery showing significant changes to the expansion of somatotopic maps and changes in electrophysiological excitability of spared motor areas post-injury (Frost et al. 2003; Eisner-Janowicz et al. 2008). Taken together, these studies provide a very concise and plausible role for spared, intact and remote cortical areas from the injury site highlighting a possible and positive role in motor recovery.

Despite these numerous neuroimaging and neurophysiological observations suggesting functional alterations occurring in these spared motor areas of the cerebral cortex, a significant vacuum exists in our knowledge and understanding of the fundamental anatomical changes accompanying these observations. To date, there are sparse numbers of non-human primate studies examining neuroanatomical changes to projection systems within spared motor areas that correlate these changes with functional levels of motor recovery (Liepert et al. 1998; Carmichael et al. 2001; Dancause et al. 2005; Stowe et al. 2007; McNeal et al. 2010). In one such study, McNeal et al. demonstrated a significant increase in the number of terminal boutons from the undamaged ipsilesional supplementary area to brachial levels of the spinal cord and correlated this increase in bouton numbers with positive levels of motor recovery (McNeal et al. 2010). However, this study only examined changes to corticofugal projection systems and did not address any cortico-cortico neuroplastic changes. However a prior, major study conducted by Dancause and colleagues in squirrel monkeys definitively showed novel and robust neuroanatomical and neuroplastic change to cortical connections existing between the distal forelimb area (DFL) of the ventral premotor cortex and the hand area of 1/2 of the somatosensory cortex at 3 months post cortical injury (Dancause et al. 2005). Indeed, both studies exhaustively demonstrated that the underlying cortical structural framework (i.e., neuroanatomical connections) can and do undergo positive neuroplastic changes.

Although this restructuring of neuroanatomical connections is exciting and opens new doors for possible interventions aimed at driving this plasticity, these studies have not properly examined nor pinpointed the early time emergence and courses of these neuroanatomical changes to cortico-cortical connections within spared motor areas and the possible correlation these changes may have in motor recovery. To address these shortcomings, we built upon the previous Dancause study and hypothesized that these new anatomical connections forming from the DFL of PMv to area occur at 4 weeks following an ischemic insult to the DFL of 1/2 3 months after an ischemic lesion to the DFL of M1. We also hypothesized that these new connections will correlate with a positive degree of motor recovery.

## Materials and Methods

All surgical exposures, cortical stimulations, lesions and injections of neural tract tracers were performed on the cortex opposite to the preferred hand as determined with behavioral analyses of reaching and flexions of the hand during initial Kluver board sessions (Nudo et al. 2003). In addition, all procedures were approved and in accordance with protocols approved by the Institutional Animal Care and Use Committee of the University of Kansas Medical Center and National Institutes of Health animal care guidelines. To accomplish the goals of this study, 6 squirrel monkeys were used in this study. Each animal underwent neurosurgical exposure of the cortex located opposite to their preferred hand as calculated using a modified kluver board (Nudo et al. 1992). Three animals were deemed controls and received a cortical map and neural tracer injections into the DFL of PMV. The other 3 monkeys also received a cortical map and tracer injections into the DFL of PMV yet had the defined M1 DFL lesioned (Table 1.).

**Table 1.** Description of Experimental Parameters in Each Case

| Case | Sex | Weight | Handedness | Tracer | Total volume<br>(nL) | Lesion Area<br>(mm<sup>2</sup>) |
| --- | --- | --- | --- | --- | --- | --- |
| 1931 | M | 660 | L | BDA | 150 | NA |
| 1926 | M | 1,100 | L | BDA | 150 | NA |
| 307A | M | 872 | R | BDA | 150 | NA |
| 735 | M | 840 | R | BDA | 150 | 9.17 |
| 830 | M | 950 | R | BDA | 150 | 14.78 |
| A42 | M | 1,100 | L | BDA | 150 | 17.38 |

### Neurosurgery

Neurosurgical exposure of the cerebral cortex of the squirrel monkey was performed under aseptic procedures and Isoflurane anesthesia as described previously (Nudo et al. 1992; Dancause et al. 2005). In brief, each animal was netted from its home cage and pre-anesthetized with ketamine HCl (100 mg/kg IM) and atropine (0.1 mg/kg IM) then transferred to the pre-surgical suite. Next, the scalp was shaved to permit an incision exposing the underlying cranium. The lower limbs were shaved below the knee for IV catheterization into the saphenous vein. A small amount of cetacaine was sprayed directly into the oropharynx to decrease any discomfort during intubation. Following the cetacaine spray, the animal was intubated with a small amount of lidocaine jelly placed on the outside of the intubation tube providing topical anesthesia and lubrication. Upon adequate intubation, each animal was moved into the main surgical suite and placed onto a ventilator and anesthetized with nitrous oxide (75 %) and Isoflurane gas (1.2-2.0%). Once an adequate of level of anesthesia was reached, an IV catheter was inserted into the saphenous vein and 3% dextrose in lactated ringer’s solution was infused along with .25cc dexamethasone. The animal was then placed into a stereotaxic head holder (David Kopf Instruments, Tujunga, CA) and a dose of 45,000 units of penicillin was given SC. The animal was attached to a life signs monitor (Digicare LifeWindow LW6000) and mannitol (~8 ml/kg IV) was subsequently administered via the IV catheter as an osmotic agent in preparation for surgical exposure of the cortex. A U-shaped incision was performed over the scalp and the temporalis muscle and underlying fascia reflected to expose the cranium and a craniotomy was performed with the underlying dura cut and reflected. A plastic cylinder was anchored in place over the craniotomy with dental acrylic and filled with warm sterile silicon oil. A small piece of a metric ruler was placed directly onto the cortex through the oil within the cylinder and a digital photograph was taken through a surgical scope (OMS 300, Topcon, Tokyo Optical Company, Japan) and transferred to a computer located in the operating room. Importantly, this photograph was used to superimpose the stimulation sites onto the 2-dimensional vascular landscape of the cortex using an illustrative program (Canvas 3.5, ACD Systems, Canada) (Fig. 2). The photograph was also used in post mortem reconstructions of stimulation maps of the DFL of M1 and the DFL of PMv (Fig. 3).

**Figure 1.**
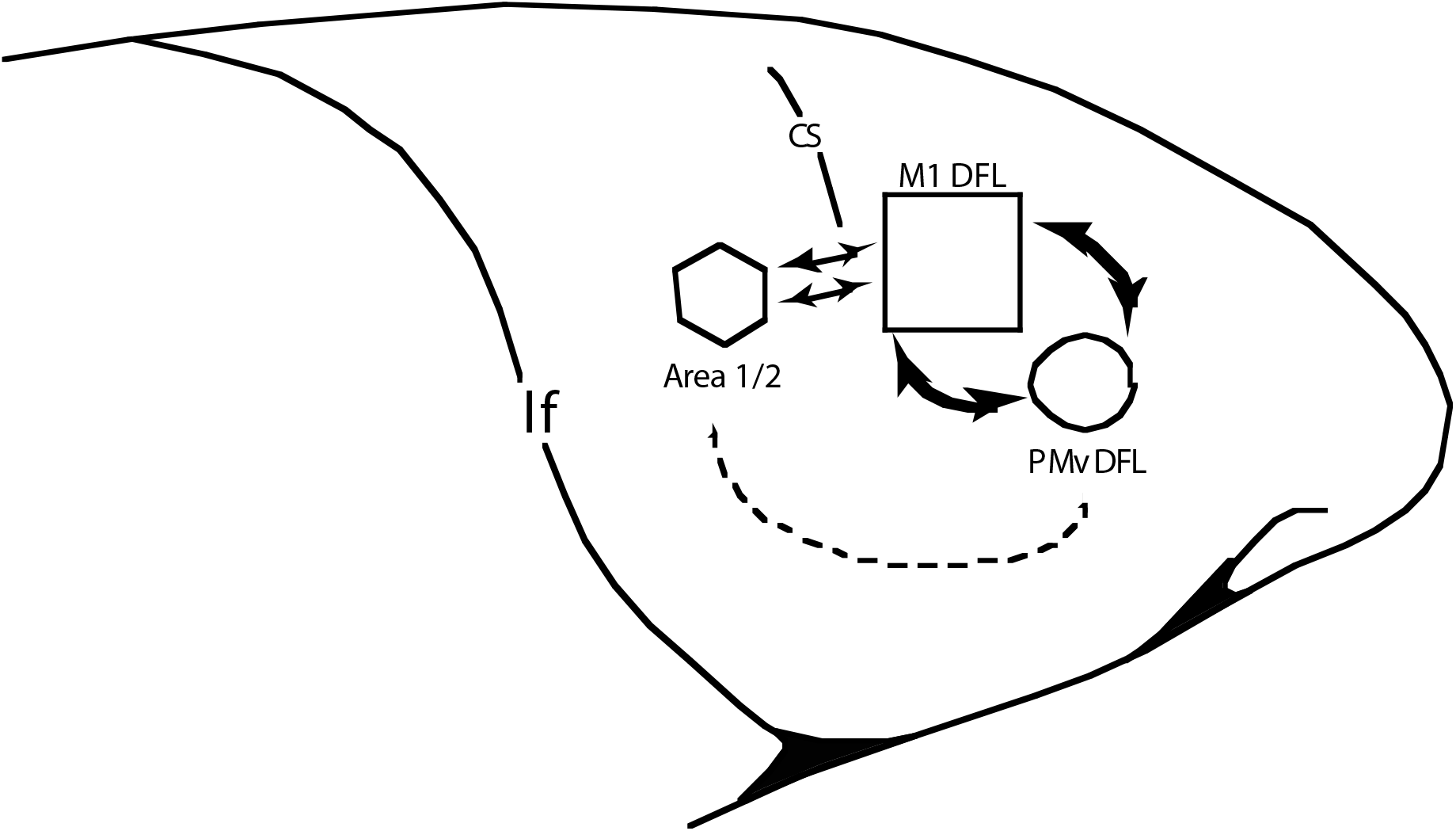
Illustration of the lateral surface of the largely lissencephalic cerebral cortex of the squirrel monkey with a focus on examining reconnections between isocortical motor areas with parietal somatosensory areas following brain injury. Of importance for our study was the examination of changes to neuroanatomical connections between the distal forelimb area (DFL) of the ventral premotor cortex (PMv) and the hand area of area 1/2 following injury to the DFL of the primary motor cortex (M1). Shown in this figure are the strong reciprocal connections between PMv and M1 and the extremely sparse connections between PMv and area 1/2. A previous study has shown an increase in the PMv to area 1/2 connection at 5 months post injury in the squirrel monkey. We hypothesized that this same plasticity would occur at a far earlier time point of 4 weeks post-injury.

**Figure 2.**
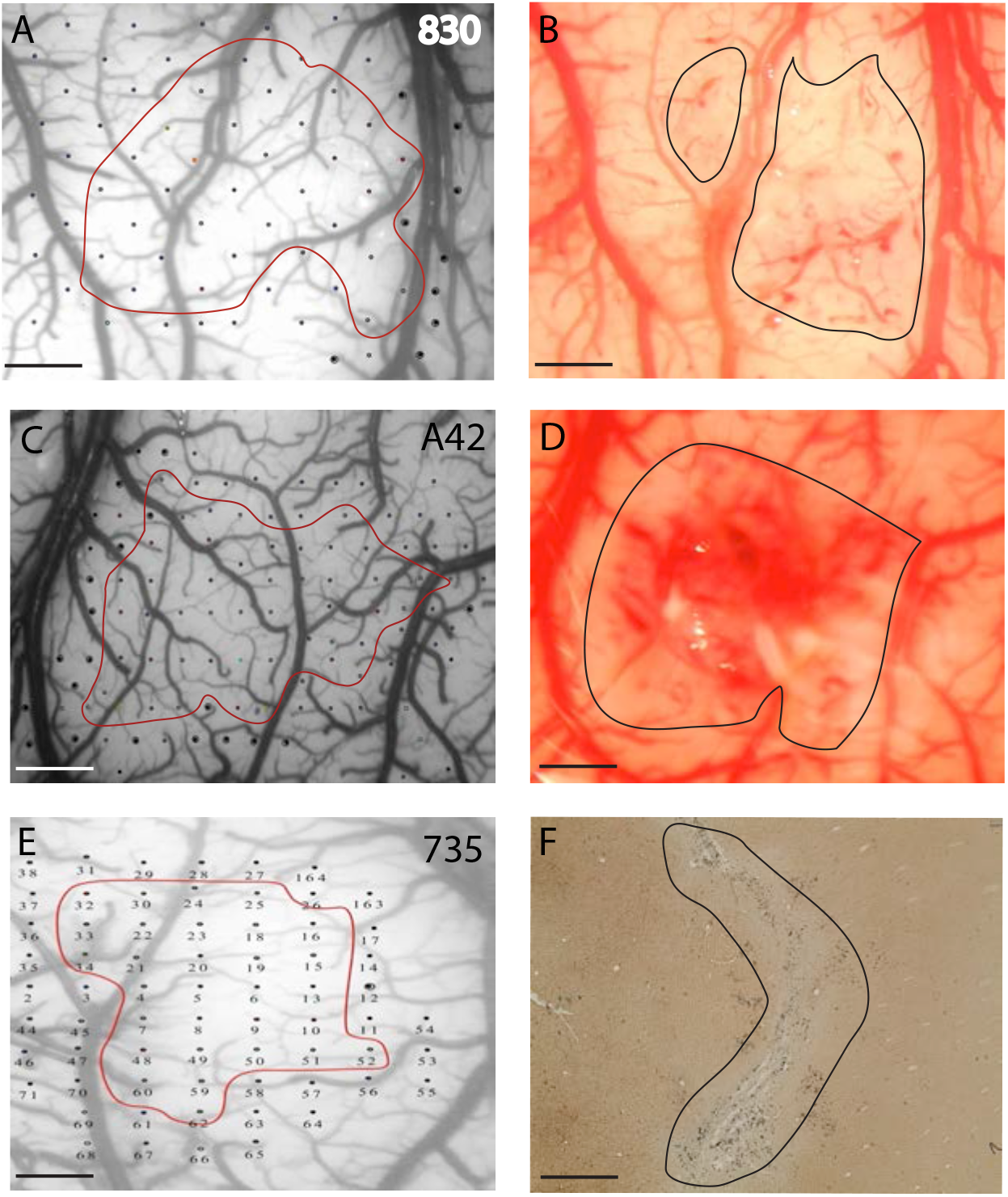
Plate of photomicrographs demonstrating the delineations of the distal forelimb (DFL) representations of the primary motor cortex (M1) and corresponding lesions (B, D and F) in cases 830 (**A**), A42 (**C**) and 735 (**E**). Stimulation points from the ICMS surgery corresponding to given body movements were superimposed on photographs of the blood vessels of the cerebral cortex of M1 (red, digits; green, wrist; blue proximal arm; yellow face; black no response). The red line delineates the DFL as determined immediately following the first ICMS surgery. The panels **B**, **D** and **F** are the defined lesions performed by electrocoagulation (black lines) and aligned across from their respective DFL stimulation maps. With the exception of case 735 whose lesion was primarily confined to caudal M1, the other two lesion cases had large lesion that affected a large percentage of the DFL of M1. Scale bars 1mm.

**Figure 3.**
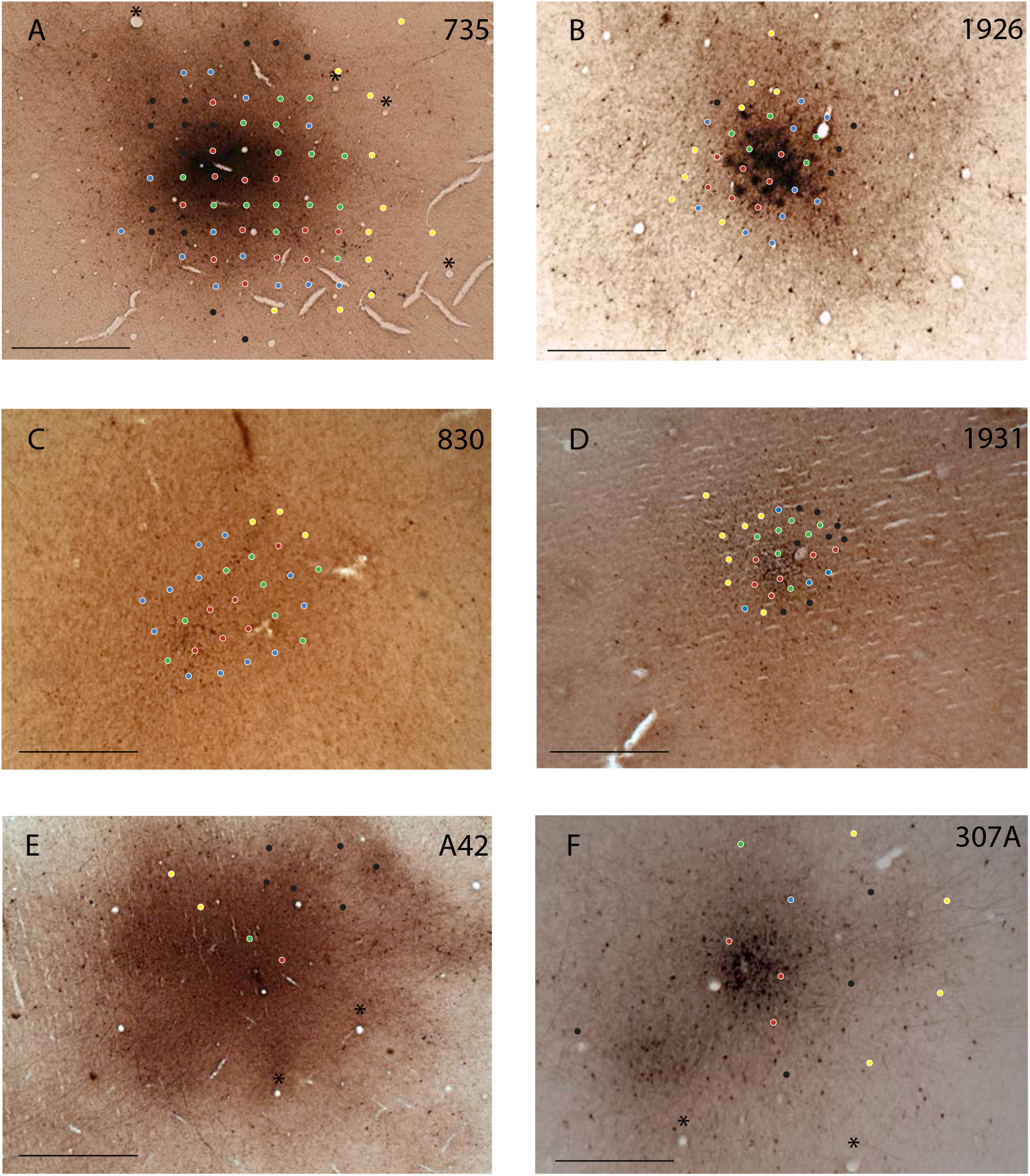
Photographic montage of Biotinylated Dextran Amine (BDA) injection sites into the DFL of PMv of lesion (cases 735, 830 and A42) and controls (1926, 1931 and 307A). The stimulation points from the ICMS surgery were coregistered with large blood vessels denoted by the asterisks on the immunohistochemically reacted tissue thereby verifying that the injection site was placed into the DFL of PMv. Each stimulation point was color-coded corresponding to a given body movement during ICMS (red, digits; green, wrist; blue, proximal arm movements; yellow, face, black, no response). Scale bars 2mm.

### Intracortical Microstimulation (ICMS) and Infarct Procedures

Prior to the electrophysiological procedures, Isoflurane and nitrous oxide gases were withdrawn and ketamine HCl (20 mg/kg IV) was injected through the IV catheter to maintain a stable anesthetic state whereby upper extremity movements via stimulation of the cortex could be elicited. Anesthetic levels were monitored using pupillary reflexes, muscle tone, heart rate and respiratory rate. A glass microelectrode (tip diameter 15-20 μm, impedance of ~600 k) filled with 3.5M NaCl and attached to a micromanipulator (David Kopf Instruments, Tujunga, Ca) was inserted perpendicular to the cortex to a depth of ~1,750 μm targeting cortical layer V. The electrical stimulus delivered was 13, 200 μs cathodal pulses delivered at 350 Hz every second from a Master 8 pulse generator (A.M.P.I.,Jerusalem, Israel) which in turn was connected to a Bak stimulus isolator (model BSI-2, Bak Electronics Inc., MD) where the amount of current delivered to the cortex could be precisely manipulated. Currents were increased until movements were evoked with a maximum current level set at 30 μA. Delineation of the distal forelimb representation (DFL) of primary motor cortex (M1) was accomplished with 500 μm interpenetration distances whereas the DFL of the ventral premotor cortex (PMv) used 500 or 350 μm spacing. Movements were recorded on the digital photograph using an illustrative program (Canvas 3.5, ACD Systems, Canada). From these photographs, reconstructions of the different cortical areas could then be analyzed.

In three control cases, we also derived somatosensory maps to define the boundary between areas 3b and 1/2 of the somatosensory (S1) cortex. To derive somatosensory maps in S1, multiunit neuronal activity were used to define cutaneous and muscle/joint fields in areas 3a, 3b, and 1/2 (Dancause et al. 2005). A Michigan-style electrode (NeuroNexus, Ann Arbor, MI) was lowered perpendicular into somatosensory cortex to span cortical layers II-V. Signals were filtered, amplified, through the RZ 5 bio amp processor from Tucker Davis Technologies (TDT, Alchulta, FL) and played over a loudspeaker for monitoring. Cutaneous responses were determined by Semmes-Weinstein monofilament 3.61 (~400 mg force) and deep receptive fields were determined by high-threshold stimulation > 3.61 and joint manipulation.

Once the determinations of the DFL of M1 and PMv were performed, the animal was placed back onto Isoflurane/ nitrous oxide anesthesia and an ischemic infarct using bipolar electrocoagulation (Malis CMC-1, Codman and Shurtleff Inc, Randolph MA, USA) of surface blood vessels was induced over the entire DFL of M1 (Fig. 2). This experimentally induced stroke method has been used repeatedly in this laboratory and has produced reliable infarcts useful for studying neuroplasticity and motor recovery (Nudo and Milliken 1996; Frost et al. 2003; Nudo et al. 2003; Dancause et al. 2006). Next, the craniotomy was closed in anatomical layers and the incision sutured. Bupivacaine was then applied to the incision (1.0 ml topical) then either sprayed with furazolidone or a topical antibacterial ointment and the animal injected with penicillin (45,000 units SC). Upon anesthetic recovery, each animal was administered acetaminophen (10-20 mg/kg oral) and codeine (1-2 mg/ml oral), then placed into an incubator and monitored every 15 minutes until stable. Each animal was then transferred back to its home cage after 10-12 hours of post-surgical recovery. Over 48 hours post-surgery, the animals were administered the same doses of acetaminophen and codeine to relieve discomfort.

Following a survival period of 2 weeks with analysis of motor recovery, a second ICMS surgery was performed upon the DFL of PMv thus enabling comparisons of changes in the DFL representation following the M1 lesion. The animals were then placed back onto Isoflurane anesthesia for injections of neural tracers into the defined DFL of PMv. Specifically, a micromanipulator was used to guide a Hamilton microsyringe (Reno NV, USA) ~ 1.75 mm below the pial surface of the cortex. Next, pressure graded injections of the tract tracer were made at depths of 1.75 mm (100 nl), 1.5 mm (50 nl) and 0.9 mm (50 nl) with a rate of 5 nl per second. Following the injections, the craniotomy was closed in anatomical layers and the animal removed from anesthesia and monitored in an incubator and given post-operative medications as described above. Motor behavior was performed following each surgery with adequate time given for the animal to recover post operatively.

### Immunohistochemistry procedures

Fourteen days after the 2nd surgery, each animal was euthanized and the cortex processed for immunohistochemical visualization of neural tracers and general cytoarchitecture according to previous procedures (Morecraft and Van Hoesen 1993; Dancause et al. 2005; McNeal et al. 2010). Each animal was immobilized with ketamine then given a large dose of euthasol IP and perfused transcardially with 3% paraformaldyhyde. Next, the brain was removed, the cortex separated from the white matter, flattened between two glass slides and placed into a 4% paraformaldyhyde/ 20% glycerol solution for 2 hrs. After 2 hrs, the cortex was placed in a 20 % glycerol/DMSO and phosphate buffer (PB) solution overnight. Next, the cortex was transferred to a 20% glycerol and PB solution overnight. The cortex was then cut on a microtome at 50 μm in a series of 3. The first series was incubated in 5% normal goat serum rinse buffer overnight then incubated in ABC solution for 4 hours then reacted with 3-3” diaminobenzidine (DAB) to visualize BDA (Veenman et al. 1992; Reiner et al. 2000) (Figs. 3,4), the second series was incubated with a monoclonal primary antibody raised in mouse directed against the neuronal marker NeuN (Chemicon, Temecula, CA) for 48 hours at a 1:1000 ratio. Following the 48 hour primary antibody incubation, tissue sections were rinsed and incubated at room temperature in a secondary biotinylated anti-mouse antibody for 4 hours (Vector Laboratories, Burlingame, CA). This series was then incubated with ABC solution and reacted with Vector SG (Vector Laboratories, Burlingame,CA) (Fig. 4). Intermittent sections from this same series were also mounted on subbed slides and stained with gold chloride to visualize myeloarchitecture (Freud, 1884) (Fig. 4). Lastly, the final series was left open to permit future studies access to lesioned tissue. All reacted tissue was then mounted on subbed slides, dried overnight then passed through graded alcohols and coverslipped.

**Figure 4.**
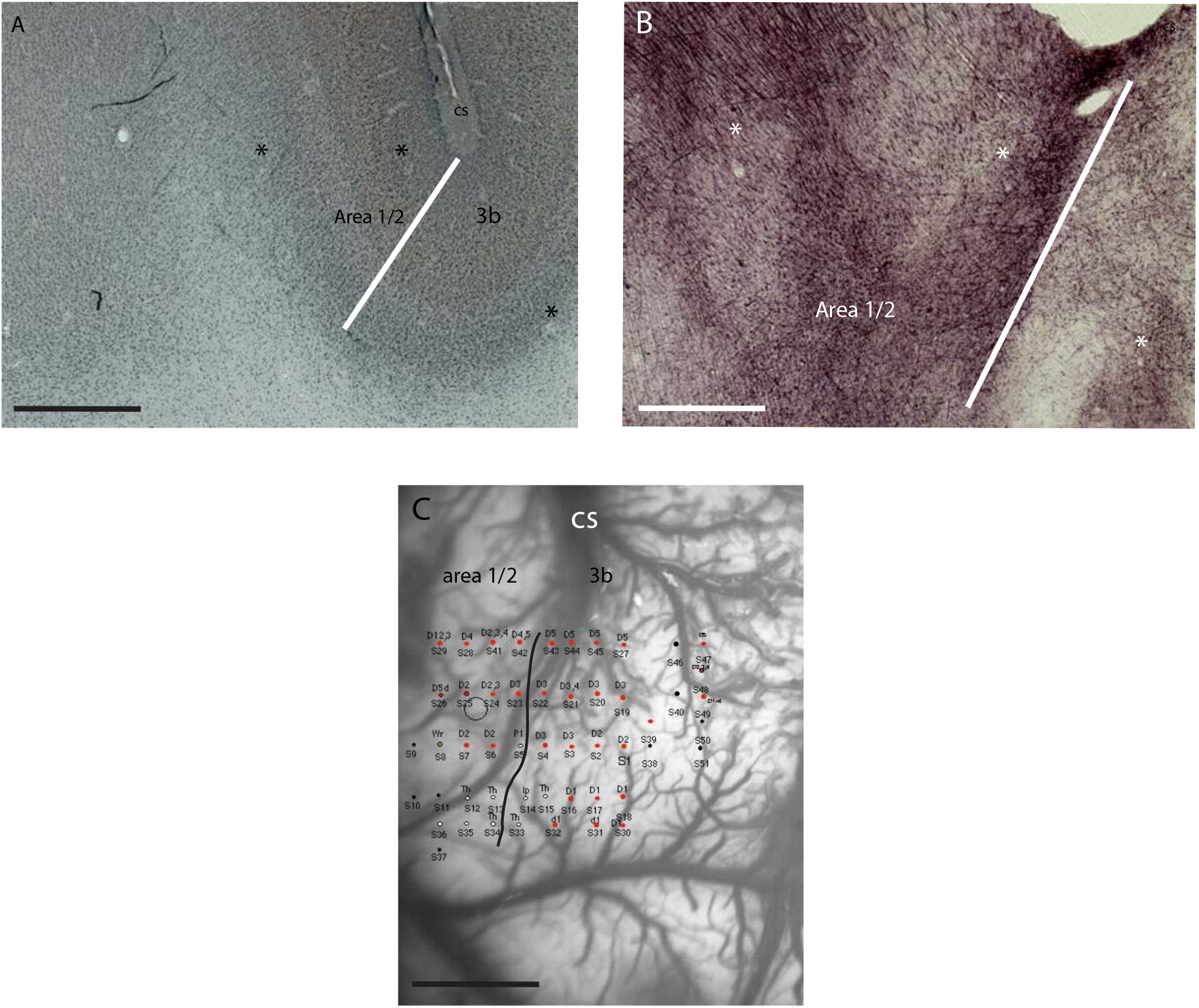
Delineation of areas 1/2 and 3b of the somatosensory cortex on horizontal sections using NeuN and gold chloride. **A**. NeuN reacted section showing Cytoarchitecture and the relative boundary between area 1/2 and 3b. We found that a defining feature of the border between these two areas was the transition from a clear layer IV and a definable layer V of 3b to a more disperse layer IV and V in area 1/2. The asterisks denote blood vessels used to align immunohistochemically reacted sections for quantification of BDA labeled terminal boutons. **B.** A gold chloride stained section staining myelin showing the border between area 1/2 and 3b. This section was spaced 100 μm from the section in **A**. Of interest is the clear demonstration of the patchwork of heavy myelin areas (dark purple) which represent digit and palm representations of area 1/2 and 3b. Asterisks denote the same vessels shown in **A**. **C.** Photograph of the lateral surface of the cortex directly lateral to the central sulcus. Recording sites were marked based upon stimulation of the hand and wrist. Digits sites were marked red and palm sites marked white. The boundary between 3b and area 1/2 was defined in a rostral caudal direction as the area where digit sites merged into palm sites and then merged into digit sites again. This physiological definition is close agreement with our histological delineation as shown in **A**. and **B**. Abbreviations: cs, central sulcus; th, thenar; d, digit; p, pad; wr, wrist.

### Data Analysis & Stereological Procedures

Analysis of immunohistochemically reacted tissue sections was accomplished under brightfield illumination on an Axioplan 2 microscope (Zeiss, Germany) with a 100x oil immersion lens (Zeiss, Germany), connected to a MAC 2000 motorized stage (Ludl Electronic Products Inc., NY). Estimations of terminal boutons from PMv were performed using the stereological probe the Optical Fractionator (Microbrightfield, Colchester, VT). This stereological probe was conducted according to methods described previously (Gundersen and Jensen 1987; Gundersen et al. 1988; West et al. 1991; McNeal et al. 2010). Briefly, the Optical Fractionator is a stereological probe that estimates population sizes by counting objects with optical disectors in a random, systematic sample within a volume of tissue located within a region of interest (ROI). Specifically, this probe uses several predetermined tissue processing parameters such as section periodicity, the region of interest area, post-reaction tissue thickness and dissector height in calculating an estimation of number of objects (see Table 2. for stereology parameters). Our target objects were immuno-reacted terminal-like varicosities that are 0.5 – 2.5 μm in diameter (Fig. 5C), and counted using unbiased counting rules (West et al. 1991). Importantly, as determined by electron microscopic analysis, varicosities of this size on the terminal region of axon fibers that are immunohistochemically identified with this type of anterograde tracer primarily represent synaptic contacts with local neurons (Wouterlood and Groenewegen 1985; Freese and Amaral 2006; Morecraft et al. 2007). Lastly, population estimates were then calculated for each animal. Next, we grouped this neuroanatomical data into control and lesion categories and compared the means. We accepted statistical significance at α= 0.05 using a two tailed unpaired t-test with a Welch’s correction.

**Table 2.** Description of Experimental Parameters in Each Case

| Case | Counting Frame<br>(W,H)<br>($\mu\text{m}$) | SRS Layout<br>(X,Y) | Top Guard<br>Zone<br>($\mu\text{m}$) | Disector<br>Height<br>($\mu\text{m}$) | Avg. Thickness<br>($\mu\text{m}$) | Series Section<br>Fraction |
| --- | --- | --- | --- | --- | --- | --- |
| 1931 | 55.81, 71.46 | 226.38, 267.55 | 3 | 12 | 18.2 | 1/4 |
| 1926 | 55.81, 71.46 | 226.38, 267.55 | 3 | 12 | 17.7 | 1/4 |
| 307A | 55.81, 71.46 | 170.84, 287.32 | 3 | 12 | 16.4 | 1/6 |
| 735 | 192.05, 71.46 | 226.38, 267.55 | 3 | 14 | 16.5 | 1/3 |
| 830 | 192.05, 71.46 | 226.38, 267.55 | 2 | 12 | 16.1 | 1/6 |
| A42 | 192.05, 71.46 | 226.38, 267.55 | 2 | 12 | 16.0 | 1/6 |

**Figure 5.**
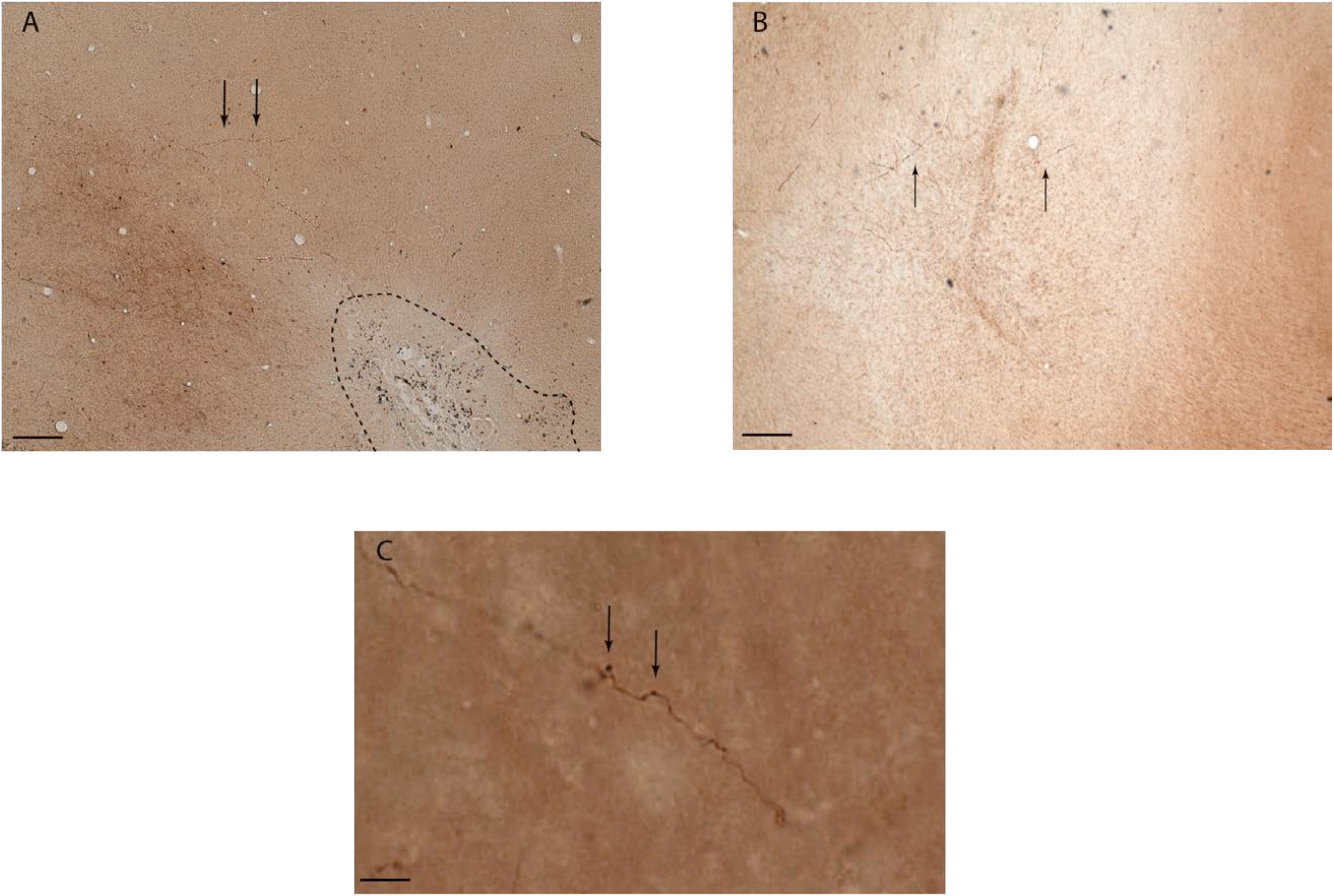
Montage of examples of BDA labeled terminal fibers and boutons. **A.** Photograph of labeled fibers from case 735 showing fibers (arrows) from PMv traveling toward the rostral part of M1 which was intact, then turning and skirting around the lesion, which is outlined with the dashed line. Scale bar is 250 μm. **B.** Labeled fibers in case A42 from PMv traveling through the glial scar that was the DFL of M1 heading towards the somatosensory cortex. Scale bar 100 μm. **C.** High magnification image of a labeled terminal fiber and a terminal bouton and an en passant bouton. This type of terminal fiber and both types of boutons were prevalent in both control and lesion cases. Scale bar 50 μm.

### Motor Recovery Testing

Assessments of pre and post-lesion motor performance were collected for each animal on a modified Kluver board (Nudo et al. 1992, 2003). Each animal’s cage was fitted with one of these boards that have five wells with varying diameters with well 1 being the biggest and in descending order well 5 being the smallest. Because of the smaller diameters, the animals must use more or less digits and or effort to retrieve a pellet from a well. This was how we scored and measured motor performance. Pre-lesion performance was used to establish baselines from which changes in motor performance can be examined. This performance was measured by the mean number of digit flexions per retrieval using the impaired limb from the Kluver board for 10 trials per 5 wells of varying difficulty. Following the M1 lesion, each animal was tested every third day post-lesion. These tests were quantified exactly as the pre-lesion trials allowing for detailed analyses of the initial motor deficit and motor recovery.

## Results

We examined the differences in terminal cortical projections from the DFL of PMv to area 1/2 of the somatosensory cortex at the time point of 4 months post-injury in 3 cases and compared the observed reorganization to 3 intact control cases. Additionally, we report positive levels of motor recovery following the cortical insult and further correlate this recovery with the degree and pattern of terminal labeling from PMv to area 1/2. However, these plastic changes occurring to the neuroanatomical connection between the DFL of PMV and the hand area of area 1/2 did not coincide with any expansion of the DFL representation as shown in previously (Dancause et al. 2005). This observation may be due to the fact we examined a relatively short survival period following the cortical injury and when compared to previous studies showing expansion of the DFL of PMV that occurred at 3-4 week post injury. As such, our study may preclude this expansion due to the short survival period following the brain injury. Importantly, our findings are the first to demonstrate this long range neuroanatomical reorganization occurring at a short survival period and will further add to add to previous studies concerning the reorganization of cortical projections of the DFL of PMv to the somatosensory cortex.

### Neurophysiological Data

Our neurophysiological cortical mapping was aimed at stimulating sites of the frontal lobe which elicited movements in order to locate and define the DFL area of M1 and PMv. Once these areas were delineated, further experimental manipulations of the cortex such as lesions and neural tracer injections were carried out. In addition, we used sensory recording to delineate areas 3b and area 1/2 in the somatosensory cortex in 3 of the 6 animals of this study. The DFL of M1 was found in an area of the frontal lobe immediately rostral to the lateral third of the central sulcus as described extensively in previous studies (Rosabal 1967; Nudo et al. 1992; Nudo 1997; Picard and Strick 2001) (Fig. 2). Within M1, the area of penetration sites eliciting DFL movements (i.e., digit flexion/extension and wrist movements) were surrounded by sites that evoked more proximal movements of the upper limb which was in complete agreement with previous studies conducting exhaustive mapping of the squirrel monkey motor cortex (i.e., elbow flexion/extension and shoulder rotations)(Welker et al. 1957; Frost et al. 2003; Dancause et al. 2006). This delineation enabled exact electrocoagulation of blood vessels and blood flow to the precise borders of the DFL of M1. In a ventral and rostral direction to the most lateral edge of the DFL of M1, we found a band of numerous sites that evoked movements of the face and even the trunk musculature. Directly adjacent to the sites eliciting face movements of M1, the DFL of PMv was characterized as a spatially distinct area of cortex that evoked distal forelimb movements and was surrounded by motor representations of cortex that evoked proximal movements in much the same way as the overall organization of M1 (Fig. 3).

We examined the cortical delineations of the hand and wrist representations of areas 3b and 1/2 of the somatosensory cortex using intra-cortical neural recordings of activity following cutaneous and deep stimulation of the hand in our control cases. We found that area 3b resided in an area of cortex directly beneath the lateral tip of the central sulcus (Fig. 4C). We found a somatotopic arrangement to area 3b that comprised digits D1 and the thenar pad laterally with a progression to D5 medially in relation to the central sulcus. Directly caudal to the digit representations, we found that the thenar pads were in close proximity to the D1 sites and that the palm and hypothenar pads were located medially. As with the maps of M1 and PMv above, our maps of the location and somatotopic arrangement of area 3b are in close agreement with other studies examining the same overall arrangement of 3b (Nudo 1997; Dancause et al. 2005; Dancause, Barbay, Frost, Plautz, Stowe, et al. 2006; Chen et al. 2011, 2012).

Immediately caudal to area 3b is the hand representation of area 1/2. This somatosensory area border was defined as place where there was a somatotopic reversal of the digits as evident by observing that in a caudal direction from the digits and palm representations of 3b, there were sites where the neural activity was induced by stimulation of the digits (Dancause et al. 2005; Dancause, Barbay, Frost, Plautz, Stowe, et al. 2006; Chen et al. 2011, 2012)(Fig. 4C). Using this reversal, we then delineated the digit representations of area 1/2 and found the same general somatotopic gradient of digit representation as in area 3b albeit in a smaller cortical area. This physiological data were confirmed with the use of NeuN (Fig. 4A) and gold chloride (Fig. 4B) to define specific borders particularly in the 3 lesion cases when there was no electrophysiological delineation of area 1/ 2 due to inadequate exposure of the somatosensory cortex.

### Lesion

Our electrocoagulation of surface blood vessels of the DFL of M1 resulted in reliable lesions that were characterized by immediate whitening of cortical areas supplied by these vessels as seen through the surgical scope. Pictures of the lesion following a period of 10-15 minutes (allowing for any incidences of reperfusion and further coagulation of these re-perfused blood vessels) were taken and loaded into the surgical computer and the area of the lesion was then calculated (Table 1.). Among the three lesion cases, we found the lesion areas to range from 9.17 mm^2^ −17.38 mm^2^ with case 735 having the most incomplete lesion (9.17 mm^2^) and case A42 having the more complete lesion of the M1 DFL (17.38 mm^2^) (Fig. 2). As per the experimental design, we attempted to occlude all of the blood vessels going into the DFL of M1. However, none of the M1 DFL areas were completely lesioned possibly owing to small vessels missed and overall re-perfusions that may have occurred following the surgery. Upon histological analysis, all three lesions spanned layers 1-6 of the cortex specifically indicating adequate damage to layers 5 and 6 which are the major locations of corticospinal neurons that terminate upon lower motor neurons located a brachial levels of the spinal cord which in turn innervate muscled of the distal forelimb, wrist and hand.

### Motor Recovery

Analyses of motor recovery before and after lesions of the DFL of the primary motor cortex were accomplished with high speed and high resolution digital motion capture cameras. For case 735, pre-lesion baseline motor performance was slightly elevated for wells 1-5 with wells 5 and 3 higher than the other 3 (Fig. 7A). Following the lesion (dotted vertical line) there was a drop in motor performance. However, this animal demonstrated a degree of scratching that the veterinarian deemed detrimental to the animal and ordered a week of doses of acetaminophen be given. This precluded our ability to administer any motor testing for post-infarct week 1. However, by post-infarct week 2 there were substantial deficits in flexions per pellet retrieval in wells 5 and 1 and by post-infarct weeks 3 and 4 motor performances returned to baseline levels. This return to baseline level was not unprecedented compared to previous studies (Nudo 1997; Dancause et al. 2005) and may be due in part to the small, crescent shape lesion located in the caudal region of the DFL of M1 that 735 received.

Animal 830 had a decent level of dexterity reflected in the modest level of stability in flexions per pellet retrieval in wells 1-4 at baseline levels (Fig. 7B). Flexions for well 5 were modestly high. Following the lesion, there was not a dramatic impact to the number of flexions per retrievals required by the animals to remove the pellets from well1-4. In contrast, well 5 had a drastic increase in the range of 4 flexions per retrieval at post-infarct week 1. This value decreased by half at post-infarct week 2 and shot up to 7 flexions per retrieval at post-infarct week 3 and finally coming down to 2.5 flexions per retrieval at post-infarct week 4 (Fig 7B). This wide range of values may be due to the fact that the lesion of 830 damaged more digits representations than 735 and since digits are required more so for well 5 retrievals, lack of these functional digits may explain these data.

Case A42 had the highest amount of variability and the highest amount of flexions per retrieval for well 1-5 for baseline levels for all 3 cases studied. Indeed, A42 had a much more complex behavioral profile (Fig. 7C). Throughout the course of the behavioral testing, this animal switched from its “preferred” hand to its “non-preferred” hand. Although this is not unprecedented (Eisner-Janowicz et al. 2008), it does require a more detailed analysis. Looking at baseline levels, A42 resembles 735, with a high flexion per retrievals of well 5 and 3 and similar 1,2 and 4 flexions. However, following the lesion, A42 switched its preferred hand and did not attempt and retrievals on wells 2-5. The animal did retrieve pellets from well 1 on post-infarct week 1 requiring 1.25 flexions slightly higher than baseline levels (Fig. 7C). However, at post-infarct week 2 there were no attempts at any of the wells. At post-infarct week 3the animal began using its affected hand again and we found that there was a high number of flexion per retrieval for well 5 and baseline levels of flexions per retrievals for wells 1-4. These data indicate that despite the animal not using the affected hand, there was an appreciable level of motor that took place with the exception of well 5.

Our behavioral data, even with several caveats demonstrate an interesting trend. In all of our cases we found that there was a positive degree of motor recovery exhibited in the form of flexions per retrieval in varying degrees of difficulty on a modified kluver board. On some of these wells, the recovery reached and improved over baseline levels.

### Injection Site Analysis

Each case in the present study received exactly 1.5 nL of BDA injectate regardless if the animal was the control group or lesion group. Our *a*nalyses of the injection of BDA within immunohistochemical reacted tissue sections for all control and lesion cases was accomplished by using the photograph and stimulation maps of the DFL of PMv acquired during the surgical procedure. The large blood vessels seen in the surgical photograph were also evident in the histological sections by small to medium size holes in the flattened cortex that retained the same spatial organization as the surgical blood vessel pattern. Using this picture, we aligned the large blood vessels seen in the photograph with the same blood vessels in the immunohistochemical sections (Fig. 3A-F). These blood vessels surrounded and were interspersed throughout the dense, dark brown precipitate located within the physiologically defined DFL of PMv. Using Neurolucida, we aligned and superimposed the reacted immunohistochemical sections with the ICMS map using these blood vessel patterns. For all of the cases in this study, the heavy, dark precipitate of the reacted BDA injectate was localized to the DFL of PMv (Fig. 3A-F). Some of the dark precipitate extended beyond the boundary where sites that elicited digits and wrist movements were located. However, this extension of injection site precipitate was minimal and a large percentage of the DFL of PMv still possessed the dark BDA precipitate allowing for the adequate examinations of neuroanatomical changes in the present study (Fig. 3A-F).

### Orientation of Fiber Trajectories

The examination of terminal labeling from the injection site was accomplished using the same flattened cortical tissue as described in our analysis of the injection site. In the control cases, we found that most fibers coursed in a caudo-medial direction to terminate within more rostral areas of the primary motor cortex (M1) in agreement with prior studies (Dancause et al. 2005; Dancause, Barbay, Frost, Plautz, Stowe, et al. 2006) (Fig. 5A). Additionally, there were a large number of fibers that traveled within the white matter in a caudo-lateral direction in relation to the central sulcus. However, within the grey matter situated directly superior to the white matter labeling, we only found few labeled fibers traveling around the tip of the central sulcus. Of particular significance for this study was our observation of the presence of labeled fibers within somatosensory area 1/2. (Fig. 5C).

In contrast to the control cases, the labeled fiber trajectories in the lesioned cases exhibited distinct changes in fiber trajectories (Fig. 5A, B). We found a fair number of labeled fibers from PMv orientating toward their normal M1 terminations yet abruptly turning lateral and traveling in a posterior direction (Fig. 5A). This change in fiber trajectory has been reported previously in the Dancause study yet we show the same changes occurring far sooner than their time point of 5 months post injury. Surprisingly, we also found evidence for labeled PMv fibers traversing through the glial scar of the DFL of M1 in case 830 and directing toward somatosensory cortex.

### Quantification of Terminal Labeling

Overall, the amount and distribution of BDA labeled boutons in intact animals from PMV to area 1/2 were similar to previous studies of this cortical connection (Dancause et al. 2005; Dancause, Barbay, Frost, Plautz, Stowe, et al. 2006) (Fig. 6A). Based upon our neurophysiological and histological delineations of the hand representation of area 1/2, we found labeled boutons throughout area 1/2 in both control and lesioned cases with a higher amount of labeling located in more caudal and lateral regions of the DFL of area 1/2. However, although these labeling trends are interesting, the focus of our study was to compare global changes in bouton numbers from PMv to area 1/2 in animals at a 4 week survival time point following an ischemic insult to M1. Thus, we did not parcelate different regions of interest within area 1/2 and quantify any bouton numbers within these separate regions. As such, we quantified the number of terminal and en passant boutons within area 1/2 as a whole in both control and lesion cases at the 4 week time interval. The estimated number of boutons for the three control cases was 4,569, 5,067 and 2,422 yielding a mean value of 4,019 (+/− 811.5 SEM) (Fig. 6B). In comparison, our lesioned animals had an estimated number of boutons of 8,806, 7,168 and 8,058 giving a mean of 9,185 (+/− 783.3 SEM) (Fig. 6B). We accepted statistical significance at α= 0.05 and using a two tailed unpaired t-test with a Welch’s correction we calculated a p value of .01 demonstrating that the difference between the means of the two groups is highly significant. The range of percentages of CEs in the control cases was (14-21%) and for the lesion cases (8-14%). The CE values represent the overall precision of the number estimation based upon our stereological sampling scheme. Importantly, the tighter range of values in the lesion cases reflects a higher number and more homogenous spacing of labeled boutons as compared to controls. Overall, for both control and lesion cases, our coefficient of error numbers of fell within acceptable levels of error and permitted us to have a high degree of confidence in our bouton estimations.

**Figure 6.**
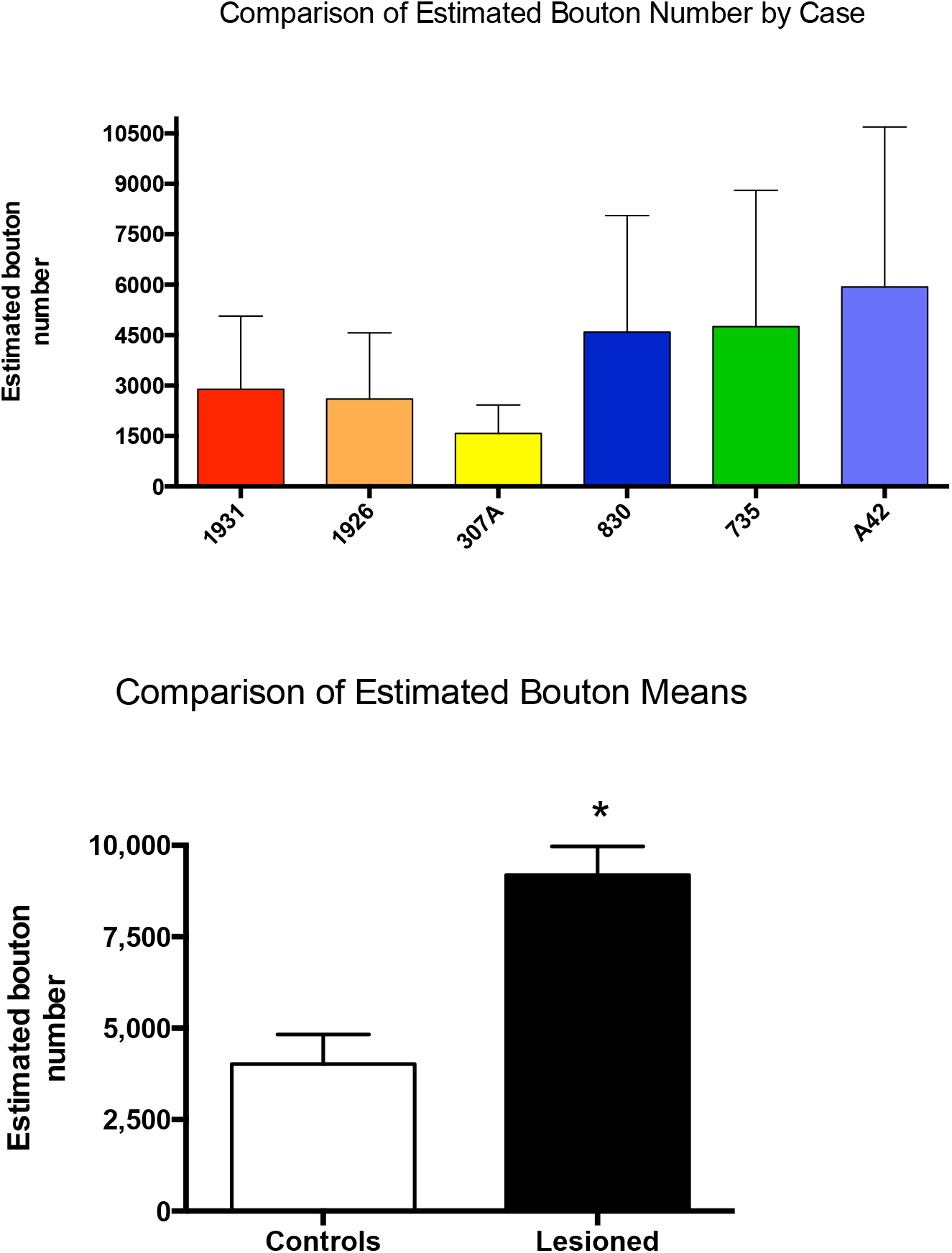
Graphs of estimated bouton numbers. **A.** Estimated bouton numbers by case. Error bars are the calculated Gundersen coefficient of errors. **B.** Grouped comparison of estimated bouton means for 3 controls and 3 lesion cases. Error bars are the SEM.

**Figure 7.**
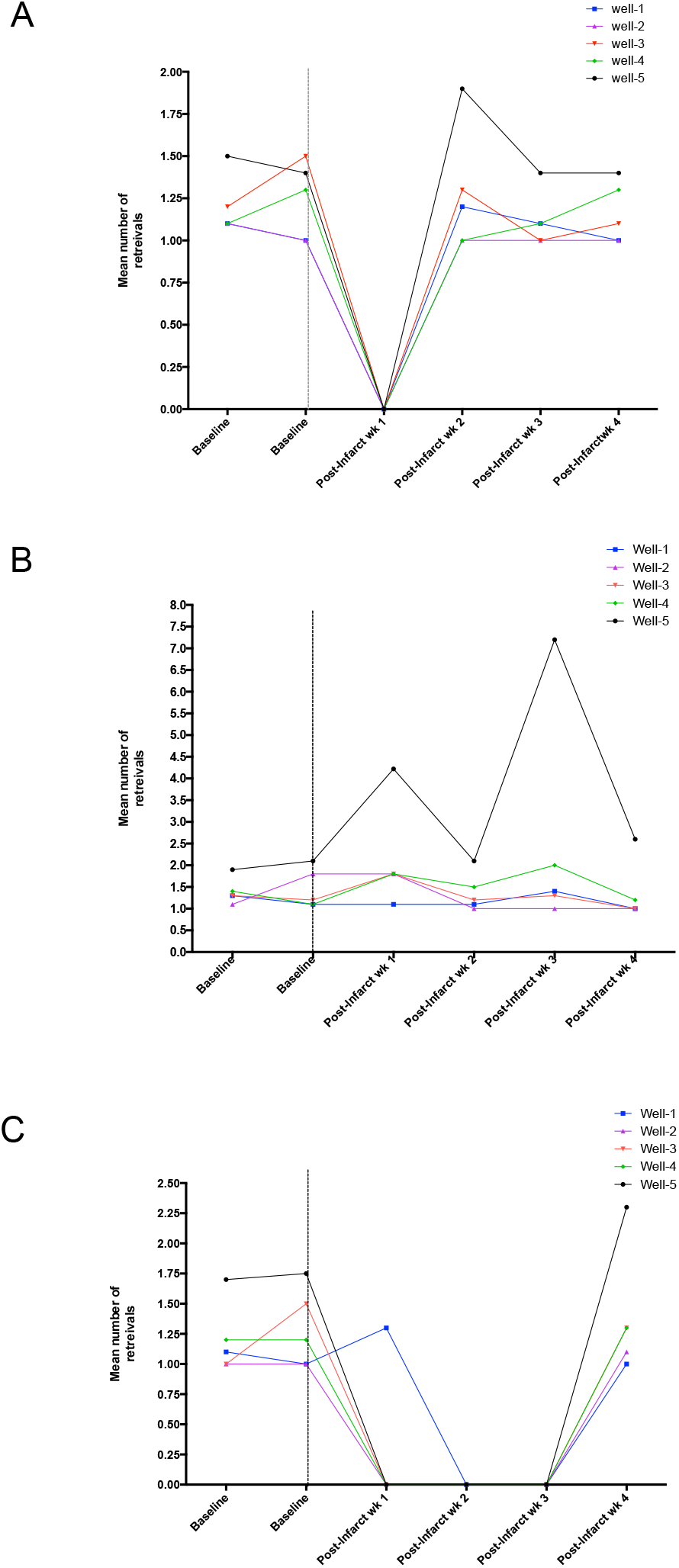
Line graphs demonstrating moderate degrees of motor recovery for all three lesion cases. **A.** Case 735. **B.** Case 830. **C.** Case A42. In all graphs the dashed vertical line represent the time of the lesion.

## Discussion

In this squirrel monkey study, we hypothesized the emergence of a robust neuroplastic response to underlying neuroanatomical connections between the DFL of PMv and area 1/2 using the anterograde tracer BDA, and that this response would occur at a ****4 week post-injury time point**** following damage to the DFL of M1. We found several important phenomena that supported our hypothesis. First, our study is the first to demonstrate a very early time point of 4 weeks post-injury for the emergence of the following neuroplastic events. Second, the neuroplasticity would manifest as altered fiber trajectories of labeled fibers from PMv that aimed to the edge of the M1 lesion and then moved laterally to enter the somatosensory cortex. In intact animals the majority of these PMv fibers would terminate directly into M1. Third, this neuroplasticity was found to nearly double the number of labeled en passant and terminal boutons from the labeled fibers from PMv to area 1/2. Importantly, we employed rigorous state of the art stereological quantification methods to arrive at a plausible estimation of these bouton numbers. To our knowledge, this study is the first to employ stereology to quantify bouton number between cortical areas in a post-injury non-human primate model. Fourth, we quantified the intact cortical projection from the DFL of PMv to area 1/2 in 3 control animals and thus provided an appreciable measure of connectivity that may be used in future studies of cortical neuroplasticity. Fifth, these post-injury changes coincided with a moderate degree of motor recovery of the affected distal forelimb. Taken together, our study reveals that PMv may be a prime area for re-establishing motor output following significant damage to M1.

### Neuroimaging and Neurophysiological Plasticity of Spared Motor Areas

For over a half a century, neuroscientific research has demonstrated the powerful ability of the brain to reorganize its physiology and neuroanatomical connections in response to significant levels of cortical injury thereby achieving some degree of lost motor function (Tsukahara 1981; Wade et al. 1985; Weiller et al. 1992; Seitz et al. 1998; Nudo et al. 2001, 2003; McNeal et al. 2010; Nudo and McNeal 2013). Indeed, this recovery has been shown to occur within the first month and extend to several years post-injury (Skilbeck et al. 1983; Wade et al. 1985; Duncan et al. 2000). However, this motor recovery is variable and not all individuals achieve similar pre-injury motor functions (Hinkle 2006; Stinear 2010). This variability in motor recovery coupled with reorganization within the cortex following injury has resulted in basic and clinical scientific inquiries concerning specific mechanisms that may be triggered and thereby play a significant role in the recovery process.

One such proposed mechanism is the role of multiple “spared”, intact and ipsilesional motor areas on motor recovery following damage to one or several of these motor areas. As each motor area possesses somatotopy, is electrically excitable and each has corticospinal projections, these additional motor areas may be large factors in determining motor recovery (Ralston and Ralston 1985; Martino and Strick 1987; Nudo and Masterton 1990; Dum and Strick 1991, 2002; Morecraft and Van Hoesen 1993; Godschalk et al. 1995; Picard and Strick 2001). Indeed, several human neuroimaging studies have shown metabolic increases in interconnected yet distant motor areas such as the Supplementary Motor Cortex and Ventral Premotor Cortex located within the ipsilesional cerebral cortex (Weiller et al. 1992, 1993; Feydy et al. 2002; Ward et al. 2003). Importantly, these researchers found modest degrees of motor recovery in patients that appeared to have metabolic increases in these motor areas. These studies shed light on the possibility that a vast reorganization of the motor network may be occurring following cortical injury to the primary motor cortex and that it may be possible these motor areas are critical in motor recovery.

In addition to the above neuroimaging studies, several neurophysiological experiments have demonstrated a reorganization of individual movements in other motor areas. For example, Frost et al. showed that at 12 weeks post M1 injury, the hand representation of PMv had a mean expansion of 36% relative to pre-injury baseline cortical map (Frost et al. 2003). In addition, another study demonstrated expansion of the DFL in the supplementary motor area at weeks 3 and 12 post injury with the expansion being directly proportional to the size of the lesion (Eisner-Janowicz et al. 2008). Furthermore, a study by Guggenmos and colleagues in rats demonstrated the effectiveness of a neural prosthesis that connected the distant locations of the cerebral cortex. They found that an injury to the caudal forelimb area disrupted links between motor and somatosensory cortices, which led to motor impairment. Once the neural prosthesis was implanted into the rostral forelimb area (RFA) and S1, spikes triggered in RFA triggered stimulation in S1 and within 1 week, the animals improved in reaching and grasping functions (Azin et al. 2010; Guggenmos et al. 2013). As a whole, these neurophysiological studies provide compelling evidence that adaptive levels of neural plasticity, in the form of changes in physiological functions, can occur in different motor areas of the frontal cortex following damage to M1 and may contribute to favorable motor recovery.

#### Relationship to Previous Studies of Neuroanatomical Plasticity of Spared Motor Areas

Expanding upon the promising neuroimaging and neurophysiological data supporting the concept of different motor areas adapting and compensating for lost cortical motor output, research aimed at investigating structural neuroanatomical plasticity has begun to offer insights into the possible role these changes may have in motor recovery. There are studies in rodents examining structural plasticity in dendritic morphological changes in response to brain injury (Brown et al. 2007, 2008) and increased synapse formation within the contralesional hemisphere (van Meer et al. 2010). However, there a very few non-human primate stroke models that have attempted to investigate neuroanatomical projection plasticity from different and distant motor areas of the brain. The importance of these studies cannot be underscored, as there are strong parallels between human and non-human primate response following brain injury studies in a non-human primate stroke model. There are however, even fewer non-human primate studies aimed at isolating time points when the emergence of this neuroanatomical plasticity occurs and peaks. Importantly, finding when this reorganization takes place will give clinicians and physical therapists a clear advantage as to when to initiate any rehabilitation.

Our study directly addresses these shortcomings of investigations into neuroanatomical plasticity by hypothesizing a single time point of 4 weeks post-injury when this plasticity will occur. Our analysis of neuroanatomical connections from PMv to area 1/2 was limited to conclusions based upon detailed analyses of the various injections sites reported in the results. We injected consistent volumes of X μL of BDA at depths of ~ 1750 μm and the areas of the injection sites between control animals and lesion cases were comparable. Thus, with a high level of confidence we conclude that the dramatic increase in bouton numbers within Area 1/2 resulting from injections of BDA into the DFL of PMv in 4 week post-injury lesion monkeys versus control monkeys was the endpoint of a massive neuroplastic response to this neuroanatomical connection.

Overall, our conclusions are in close agreement with a previous study that found a significant increase in bouton number from the DFL of PMv to the area 1/2 hand area in the squirrel monkey at 3-5 months post M1 DFL injury (Dancause et al. 2005). Importantly, Dancause and colleagues also demonstrated that a moderate degree of motor recovery coincided with this neuroanatomical reorganization. Interestingly, their study found no significant changes to connections between other somatosensory cortical areas such as areas 3a and 3b which have strong connections to M1. As our study only built upon examining the significant projection of PMv to area 1/2 we cannot directly compare to this aspect of their study which involved several different cortical areas. However, besides the examination of a four week time point in our study, we employed rigorous stereological quantification of terminal and en passant boutons within area 1/2. The Dancause study used a voxel based approach to their quantification where a marker is placed on a grid that overlays the cortical surface. Different markers corresponding to the number of boutons observed were then placed into the grid square. Although this method is an excellent approach to mapping out cortical connections, it does not provide a solid number of boutons that would indicate and appreciable level of connectivity. Our study provides real estimations of the number of boutons and thereby permits more robust statistical descriptions and inferences. However, future studies may capitalize on examining other cortical motor and somatosensory areas in terms of neuroplasticity and subsequent motor recovery and use the findings of the Dancause study as a point of reference.

#### Clinical Relevance and Importance

As expanded upon above, several rodent and non-human primate studies have posited the possible functions and underlying mechanisms that both contralesional and spared ipsilesional motor cortices may have on motor recovery. What are lacking in these studies is an adequate time point of when these mechanisms and or functions may occur post-injury and the application of rigorous stereological quantification of either synaptic contacts or terminal fiber length. By being the first to design our experiment and to hypothesize increases in bouton numbers between the DFL of PMv and area 1/2 at a 4 week post-injury endpoint and the use of stereological quantification of boutons from the DFL of PMV to area 1/2 we have addressed these shortcoming in previous studies and have provided novel insight into the neuroplastic response and the role that area 1/2 may play in this function following subtotal brain injury. The two cortical areas we focused upon are major motor and somatosensory cortical centers that are critical in motor recovery particularly at the clinically relevant 4 week post-injury time point. This reorganization and re-establishing of connections between motor areas and sensory areas may be in fact be an intrinsic property of the damaged brain to re-establish a degree of somatomotor control (Roiha et al. 2011; Laaksonen et al. 2012). Indeed, this time week point of 4 weeks post-injury may allow for resolution of diaschisis events through the motor network and may occur prior to the formation of a dense glial scar that may prevent any long-term restructuring of neuroanatomical connections between PMv and area 1/2.

Also, several studies have shown that reorganization of movement maps as defined by ICMS occurs 1-6 weeks following the cortical injury (Eisner-Janowicz et al. 2008). Although there may exist a temporal mismatch between map changes and behavioral recovery, there exists the possibility that the neuroanatomical reorganization may undergo a longer period of manifestation due to molecular events and the overall distance traveled. Additionally, this 4 week time period is critical as many clinicians and therapists point to and stress early intervention to better promote the motor recovery process. As such, because of our finding of a 4 week plastic response, clinicians may be able to capitalize on the inherent properties of the cerebral cortex in response to injury and combine these properties with therapeutically designed ones.

Of high importance, in all three cases we found favorable levels of motor recovery with the majority of them achieving baseline levels with the exception of well 5 which is the toughest well. This lack of retrieval of pellets from well 5 may show that the animals did not recovery completely, but it is important that these animals did not undergo and rehabilitative training and thereby classified as “spontaneous” recovery as opposed to what would be conducted in human cases.

Our findings in this squirrel monkey study are relevant in humans suffering from sub-total brain injury in several ways. There are homologous motor areas in both monkey and human that may possess the same process for neuroplastic reorganization following brain injury. Also, each of the motor areas possess a somatotopic arrangement, each motor area is highly and reciprocally interconnected with several somatosensory areas such as 5 and 7 and PO/PI, all motor areas are electrically excitable, and each predominantly control musculature opposite from a given hemisphere via the spinal cord (Ralston and Ralston 1985; Martino and Strick 1987; Nudo and Masterton 1990; Dum and Strick 1991; Morecraft and Van Hoesen 1993; Godschalk et al. 1995). However, it should be noted that the main function that has been found in the literature concerning the function of PMv is that it has a strong facilitating action upon M1 by directing movement initiation and patterns and also may integrate proprioceptive and fine sensitive sensory functions (Kakei et al. 2001; Cerri et al. 2003; Shimazu et al. 2004). Despite this strong action on M1, the main corticospinal terminations predominantly innervate cervical levels 1-4 with a small percentage innervating cervical spinal cord level five (Martino and Strick 1987; Nudo and Masterton 1990; Dum and Strick 1991). Hence, PMv does not possess the direct connections to lower levels of the brachial spinal cord which directly influence and control distal forelimb muscles. As such, PMv may require numerous connections both cortical and sub-cortical to influence motor output.

Another observations that we found in our study concerned the concept of a lack of sensory feedback from the affected hand, which is also a common occurrence in human patients suffering from stroke (Carey et al. 1993). Our lesioned animals with large deficits presented with classic signs of a form of sensory neglect in the affected hand that may further impair their motor performance as mentioned above. That is, the animal may retrieve the pellet, which is a motor function, yet that animal does not realize these is a pellet in the hand until visually noticing it. This fact reflects that not all lesions affect only motor structures and underscores the importance of re-establishing a connection between a heavily represented hand area of PMv with a hand area of area 1/2. Indeed, this strengthening of connections between PMv and area 1/2 may become a potent modulator for positive levels of motor recovery.

Within our neuroanatomical data, we found that the majority of boutons located in area 1/2 were located in more ventral aspects closer to ventral fasciculi. This finding may be important as the connection from PMv to posterior levels of the parietal lobe follow these inferior fasciculi (Sakata and Taira 1994; Luppino et al. 1999). However, these previous studies do not in any way diminish ours and previous findings of labeled PMv fibers actually skirting and re-directing around the M1 lesion on their course to area 1/2 (Dancause et al. 2005) (Fig. 5A).

### Alternative Explanations and Technical Considerations

#### Other Motor Areas and Other Connectional Pathways

The current study focused only on neuroplasticity of horizontal connections within the cerebral cortex. Although this study is the first to employ rigorous stereological quantification of the plastic changes to these neuroanatomical changes between the DFL of PMV and area 1/2 at a clinically relevant post-injury time point of 4 weeks point injury, However, there are studies that have demonstrated that this plasticity is not solely the domain of PMv cortical connectivity. As there are several individual motor fields located in the cortex and each of these motor fields has corticospinal connections, there exists the possibility that these motor areas may assist or even take over the lost motor functions after brain injury with their own neuroanatomical plasticity prior to or in conjunction with PMv neuroanatomical plasticity. In one major non-human primates study, McNeal and colleagues demonstrated that the corticospinal projection from the ipsilesional Supplementary Motor Area undergoes a high degree of neuroplastic bouton proliferation at brachial levels (C5-T1) of the spinal cord at 6 months and 1 years following surgical resection of the arm and hand representation of M1 and the dorsal lateral premotor cortex (McNeal et al. 2010). Thus, it is clear that the entire motor network may be contributing to reestablishing a degree of previous motor output and we cannot solely focus on one motor area over the other. Instead, the much more more prudent and effective approach will be to employ holistic therapies aimed a capitalizing on all motor areas and there similar and sometimes unique outputs.

While most of these other spared motor areas have been ipsilesional and their cortical connections have been ipsilesional as well, recent interest has revealed that contralesional motor fields and their corticospinal sytems may indeed play a large role in the motor recovery process. Lindau and colleagues have found that rats that received a unilateral photothrombotic stroke that damaged greater than 90% of the sensorimotor cortex improved their skilled reaching to 65% of that of pre-lesion (Lindau et al. 2013). Upon histological examination, these researchers used anterograde tracers from the contralesional cortex and found a 2-3 fold increase in in midline crossing by neutralizing the CNS growth inhibitory protein Nogo-A. This is surprising as the corticospinal tract maintains nearly a 95% uncrossed trajectory. They concluded that this crossing led to the functionality of the motor behavior following the injury and provide strong evidence for the role of the contralesional cortex in motor recovery after brain injury.

A major technical aspect concerns our description of BDA filled varicosities (boutons) as putative synapses. AS determined by several electron microscopic studies, varicosities of the size we counted (.5μm-2.5μm) located on either the terminal branches (terminal boutons) or the mid-levels of the axons themselves (en passant) which are immunohistochemically identified using state-of-the-art anterograde tracers, primarily represent synaptic contacts with neuron profiles (Wouterlood and Groenewegen 1985; Freese and Amaral 2006). As such, our goal of this study was not to investigate the ultrastructural anatomical organization of these boutons but to estimate a degree of axonal connections to a large area of somatosensory cortex in the squirrel monkey cerebral cortex. We attempted to complete this by using these terminal-like profiles offered by highly sensitive anterograde tract tracing and immunohistochemistry. However, we cannot rule out the possibility that some of these boutons that we quantified may indeed not be synaptic contacts and may be merely axonal swelling. AS it is beyond the scope of this paper, future studies will need to be designed and employed to directly address this technical detail.

The previous study, upon which this case was modeled upon (Dancause et al. 2005), employed the use of the Gallyas myelin stain to delineate the borders between areas 3b and area 1/2. Furthermore, this stain beautifully showed the digits the face/hand border of 3b and the digits of the hand area of area 1/2. As such, the Dancause study was easily able to quantify voxels within area 1/2 and compare control and lesion cases. In our study, we extensive cortical sensory maps of areas 3b and area 1/2 where we are able to show the distinction between 3b and area 1/2 based upon a flip in the somatotopy. Unfortunately, in all 3 of our lesion cases, the surgical exposure and difficulties during the surgical procedure precluded us from being able to perform a sensory map. Thus, our only recourse was to use post mortem cytoarchitecture and myeloarchitecture and blood vessel patterns to delineate areas 3b and area 1/2. We are confident that we have performed this adequately, but we must note the difference in reconstructions between our control cases and experimental and between the Dancause study.

We have shown that between our controls and experimental animals there is a robust neuroanatomical plastic response that occurs at 4 weeks post-injury to the DFL of M1. Importantly, we show that there is a more than moderate degree of motor recovery that occurs in all three experimental animals with some of them even performing better than baseline on certain wells. We must assume that we can draw no definitive causation between this neuroplastic response and the motor recovery. One can only speculate that it is correlative as we cannot posit any solid mechanistic hypothesis as to what is driving this motor recovery. True, the neuroplastic response may play a large role, but there may be other factors that may play the same large role or an even larger role. As such, we can only take the stand that our study appears to positively correlate with positive levels of motor recovery.

## Summary and Conclusions

This study is the first to demonstrate a robust neuroanatomical neuroplastic response occurring to major horizontal connections of the cerebral cortex in the non-human brain following a four week post-injury survival period. This plasticity took the form of dramatic increases in bouton numbers, as quantified stereologically, between the DFL of PMv of the premotor cortex and the hand representation of area 1/2 of the somatosensory cortex. In agreement with other studies, we hypothesized that the neuroplastic response we observed was an effort of the cerebral cortex to re-establish equilibrium between somatosensory input to motor areas and novel motor output points thereby allowing adequate functions of daily behavior and living. Building upon this hypothesis, this neuroplastic response was accompanied with a moderate degree of motor recovery, which indicated that our observed 4 week plasticity may indeed play a large role in decreasing the time of motor recovery following sub-total brain injury.

